# Soil organisms in military-impacted environments: A systematic review of microbial community studies, contamination types, and methodological gaps

**DOI:** 10.64898/2026.03.13.711562

**Authors:** Naomi Beddoe, Naomi Rintoul-Hynes

## Abstract

Military activities can significantly influence soil ecosystems through physical disturbance and the introduction of contaminants such as explosive compounds, heavy metals, and hydrocarbons. Soil microbial communities play key roles in ecosystem functioning and contaminant transformation, yet the extent to which these communities have been systematically studied in military-impacted soils remains unclear. This study presents a systematic review of research investigating soil biological communities in landscapes affected by military activity or warfare.

A structured literature search was conducted across Scopus, Web of Science, Dimensions, PubMed, and Google Scholar. Following duplicate removal and multi-stage screening, 20 studies met the inclusion criteria. Data were extracted on study location, contamination types, soil physicochemical measurements, biological methods, and methodological characteristics. A Methodological Completeness Index (MCI) was calculated to evaluate the extent to which studies integrated environmental and biological measurements.

Results reveal a strong reliance on 16S rRNA amplicon sequencing, used in 80% of studies, while fungal and soil fauna investigations were rare. Soil physicochemical characterization was inconsistent: soil pH was measured in 60% of studies, whereas microbial biomass and enzyme activity were reported in fewer than 20%. No studies reported soil bulk density despite the importance of soil compaction in military landscapes. Research focused mainly on explosive compounds and heavy metals, particularly TNT, RDX, and lead contamination.

## 1 Introduction

Military activities have long been recognised as significant drivers of environmental change across terrestrial ecosystems (Certini & Scalenghe, 2013). Training exercises, weapons testing, ammunition disposal, and armed conflict can alter soil systems through a combination of physical disturbance, chemical contamination, and landscape modification (Broomandi et al., 2020). Activities such as vehicle movement, artillery impacts, and explosive detonations may cause soil compaction, crater formation, and structural disruption, while military infrastructure and operational activities can introduce a range of contaminants including explosive compounds, heavy metals, hydrocarbons, and chemical warfare residues (Prost, et al., 2013; Broomandi et al., 2020). These disturbances can have profound effects on soil properties and ecological processes, potentially altering microbial community composition and ecosystem functioning (Rodríguez-Seijo et al., 2024).

Soil microbial communities play a central role in maintaining soil health and ecosystem resilience. Bacteria, fungi, and soil fauna contribute to processes such as organic matter decomposition, nutrient transformation, pollutant degradation, and soil structure formation (Philippot et al., 2024; Wang et al., 2024). These microbial processes regulate biogeochemical cycling and strongly influence soil ecosystem functioning. Changes in soil biological communities can provide valuable insights into the ecological impacts of disturbance and contamination. Advances in molecular techniques, particularly high-throughput sequencing and metagenomics, have enabled increasingly detailed investigation of soil microbial diversity and function Garg, 2024); Reznikova, et al., 2025). These approaches allow researchers to analyse microbial genetic material directly from environmental samples, providing insights into microbial community composition and functional potential. Such methods have been widely applied under various land uses including agricultural (Wilhelm et al., 2023), forest (Chan et al., 2006), and contaminated soils, although their application in military-impacted landscapes remains comparatively limited.

A growing body of research has begun to examine microbial communities in soils affected by military activities. This includes explosive compounds such as trinitrotoluene (TNT), hexahydro-1,3,5-trinitro-1,3,5-triazine (RDX), and octahydro-1,3,5,7-tetranitro-1,3,5,7-tetrazocine (HMX) (Lewis et al., 2004; Corredor et al., 2024). These compounds are known to influence microbial activity and may be degraded or transformed by specific microbial taxa (Corredor, et al., 2024; Yang, et al., 2021). In addition to explosives, military soils may contain heavy metals and metalloids associated with ammunition residues, including lead, copper, zinc, and antimony, Other contaminating agents include hydrocarbons from fuel spills. These explosives, heavy metals, metalloids and other agents can contaminate the soil through warfare, training and other operational activities including (ref, date). Despite increasing interest in these environments, studies vary widely in their methodological approaches, environmental measurements, and geographic focus.

Understanding how military activities affect soil biological communities requires an integrated assessment of contamination, soil physicochemical properties, and microbial community structure. However, it remains unclear to what extent existing studies provide comprehensive environmental characterization or whether research efforts are concentrated on particular contaminants, organisms, or geographic regions. A systematic synthesis of the literature is therefore needed to identify current research trends, methodological gaps, and priorities for future investigation.

The aim of this study was to conduct a systematic review of research investigating soil organisms in military-impacted environments using sequencing-based and related community profiling approaches. Specifically, this review sought to:

- Identify studies examining soil biological communities using sequencing-based and related community profiling approaches in landscapes affected by military activity or warfare.
- Evaluate the biological methods used to characterise soil organisms.
- Assess the soil physicochemical and contamination variables measured in these studies.
- Examine the geographic distribution of research efforts.
- Evaluate methodological completeness using a Methodological Completeness Index (MCI).

By synthesizing available evidence, this review aims to provide a clearer understanding of how soil organisms are studied in military-impacted environments and to highlight key gaps that should be addressed in future research. Figure 1 provides a conceptual visual representation of military activities, contamination categories and what those activities and contamination agents may affect in the soil in terms of soil properties and soil organisms.

**Figure 1.**
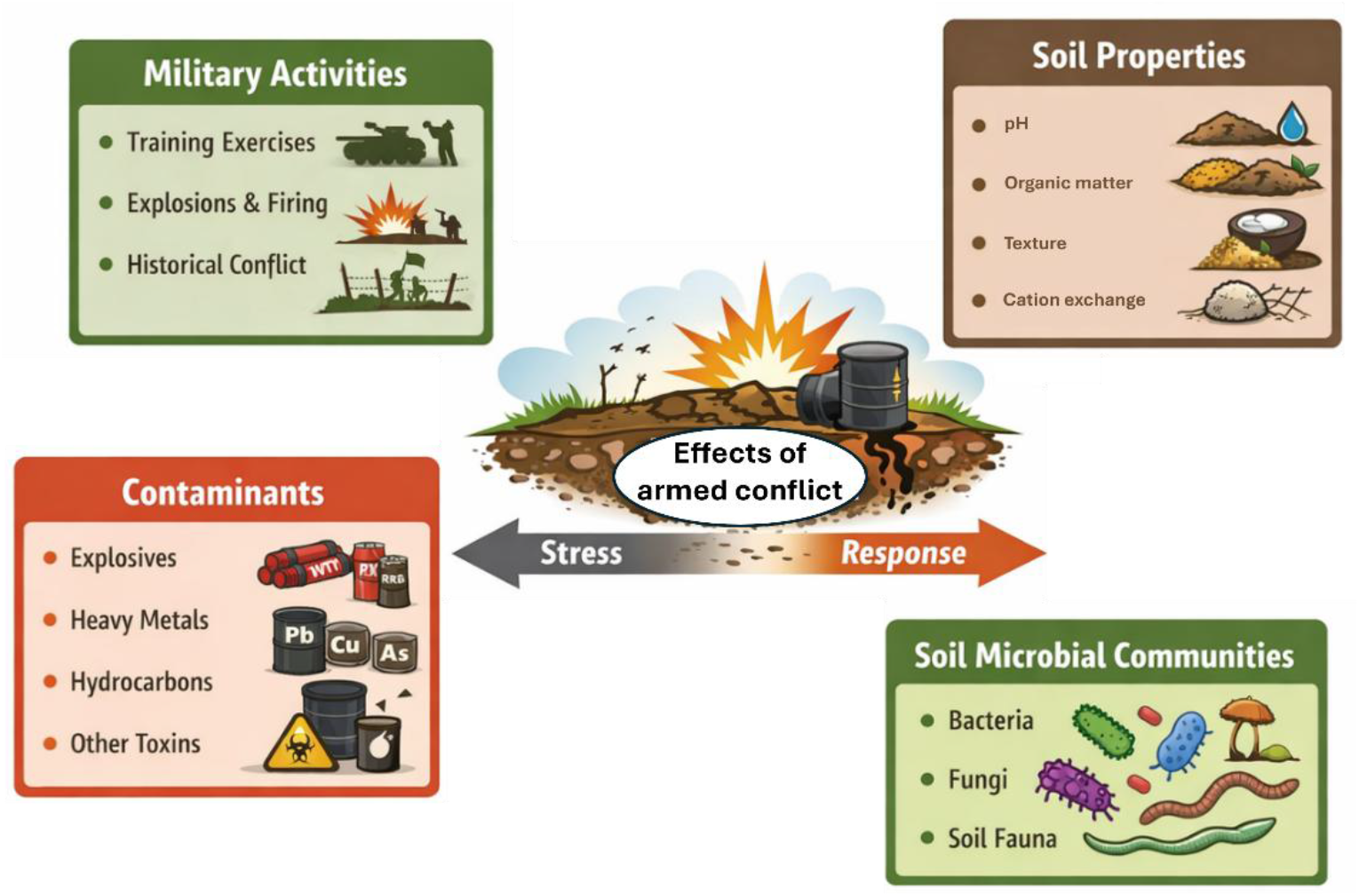
Conceptual framework illustrating the interactions between military activities, contaminant inputs, soil physicochemical properties, and soil biological communities. Military activities such as training exercises, weapons firing, and historical conflict can generate soil disturbance and introduce contaminants including explosives, heavy metals, hydrocarbons, and other toxic compounds. These disturbances alter key soil physicochemical properties, including pH, nutrient availability, organic matter content, moisture, and cation exchange capacity. Changes in soil conditions influence the structure and function of soil biological communities, including bacteria, fungi, and soil fauna. In turn, microbial communities can mediate contaminant degradation, transformation, and ecological recovery processes. The framework highlights the interconnected pathways through which military disturbance and contamination influence soil ecosystem functioning. This framework summarizes the environmental context investigated across the studies included in this systematic review.

## 2 Methods

### 2.1 Database search strategy

A structured literature search was conducted to identify studies investigating soil organisms and microbial communities in environments impacted by military activities or warfare. Because terminology used in this field spans multiple disciplines including soil science, environmental microbiology, and contamination research, the search strategy was designed using functional keyword blocks representing four conceptual components: (1) soil or sediment environments, (2) soil organisms or microbial communities, (3) biological community profiling methods, and (4) military or conflict-related site descriptors. This approach allowed the search strategy to capture studies describing both contemporary molecular analyses and earlier microbial community profiling methods while maintaining sufficient specificity to identify research conducted in military-affected environments.

Databases Scopus, Web of Science, Dimensions PubMed and the search engine Google Scholar were used for identifying studies for inclusion in the review. Filters were used to filter returned items with a data range of 2000-2026, the English language, and for article publication type.

Prior to finalizing the database search strategies, an exploratory analysis of terminology used in the pilot dataset was conducted. Titles, abstracts, and keywords from previously identified records were analysed using a Python-based text mining approach to identify frequently occurring words and phrases related to soil organisms, community profiling methods, military activities, and landscape disturbance. The resulting term frequencies were used to refine the functional keyword blocks applied in the final Boolean search strategies, helping to ensure that the search queries captured relevant variations in terminology used across the literature

#### Scopus

Scopus was searched using a structured Boolean strategy applied to the title, abstract, and keyword fields. The search combined functional blocks representing soil or sediment environments, soil organisms or microbial communities, biological community profiling methods, and military or conflict-related site descriptors. Additional terms were included to capture both modern sequencing approaches and earlier community profiling techniques. Searches were limited to publications from 2000 to 2026, and results were exported for subsequent deduplication and screening.

#### Web of Science

Web of Science Core Collection was searched using the Topic field, which indexes titles, abstracts, author keywords, and Keywords Plus. The search strategy used the same conceptual blocks as the Scopus search but was adapted to Web of Science syntax and interface constraints. Filters were subsequently applied for English-language articles and the publication period 2000 to 2026. The resulting records were exported in spreadsheet format for inclusion in the screening pipeline.

#### Dimensions

Dimensions was searched using a structured keyword strategy combining soil, microbial community, method, and military-context terms. Because Dimensions indexes broad searchable record content and has limited support for field-restricted Boolean logic, the query was adapted to improve precision while retaining recall. Searches were restricted to the period 2000–2026, and results were exported for downstream deduplication and screening.

#### PubMed

PubMed was searched using field-restricted terms applied to the Title/Abstract fields. The query combined soil or sediment terms, microbial community and organism terms, sequencing and profiling methodology terms, and military or ordnance-related descriptors. Filters were applied for English-language articles published between 2000 and 2026. PubMed was included to capture microbiology-focused studies that may not have been indexed consistently in broader environmental databases.

#### Google Scholar

Google Scholar was used as a supplementary search source to identify potentially relevant records not captured in the structured database searches. Because Google Scholar does not support controlled export or highly reproducible field-specific filtering, targeted keyword combinations were used and results were screened manually. The first results returned by each query were examined, and relevant studies were recorded in a structured spreadsheet for integration into the review workflow. This stage was used primarily to improve search completeness and identify additional records from less well-indexed sources.

### 2.2 Study identification and deduplication

Studies were identified by searching database/search engine as previously described (2.1).Returned filters items were exported to CSV/Excel. Consolidation of records, duplicate detection and removal (85 records) by DOI was achieved by a Python script.

### 2.3 Eligibility criteria

#### Inclusion criteria

- soil or sediment samples
- military-impacted environments
- contamination relevant to military activity
- biological community analysis
- sequencing or community profiling methods

#### Exclusion criteria

- non-soil environments
- studies without biological community analysis
- non-military contamination sources

### 2.4 Screening workflow

Stage 1 screening was automated using a Python script with keyword blocks. This output categories of likely include, manual review and likely exclude. Stage 2 screening was manual for the likely include, manual review and 127 of the likely exclude returned studies. The manual element of the stage 2 process was verified by two additional researchers, who also manually screened the likely include, manual review and 127 of the likely exclude returned studies, blind from the first reviewers results. PRISMA compliance numbers were recorded. Several additional studies from earlier searches were included and accounted for, giving the final included record pool. Following full text reading it was decide that 3 records were then to excluded as they lacked explicit military linkage. To support the initial screening process, a set of custom Python scripts was developed to assist with record processing and prioritization. These scripts standardized exported records from different databases, merged datasets, removed duplicate entries using DOI identifiers, and applied rule-based text scanning to titles, abstracts, and keywords. The automated screening stage used predefined keyword patterns representing soil environments, microbial communities, biological profiling methods, and military-related site descriptors to assign preliminary relevance classifications (e.g., likely include, manual review, or likely exclude). This approach allowed rapid prioritization of records while ensuring that final inclusion decisions were confirmed through manual review. The automated workflow was used solely to support and accelerate screening, and all records classified as potentially relevant were assessed manually to confirm eligibility. The automated workflow facilitated efficient processing of the >6,000 records identified across multiple databases. The review followed the Preferred Reporting Items for Systematic Reviews and Meta-Analyses (PRISMA) guidelines.

### 2.5 Data extraction

Data extraction was performed manually using a standardised Excel spreadsheet template. The variable categories extracted were study metadata, site characteristics, soil physicochemical variables, contaminants measured, biological methods and organism groups.

### 2.6 Data synthesis and analysis

The standardised Excel spreadsheet template collated study metadata including Study ID, First author, year, article title, journal name, digital object identifier (DOI) and study type. It collated information on site type including country, region, latitude, longitude, military site type, current or historical military activity and landscape type. It counted the frequency of soil physiochemical measurements (pH, soil organic carbon (SOC), soil organic matter (SOM), total nitrogen (TN), the carbon nitrogen ration (C/N or C:N), soil texture, sand/silt/clay percentage, soil moisture, bulk density, electrical conductivity (EC), cation exchange capacity (CEC), enzyme activity and microbial biomass). It counted the frequency of records that studied explosives and specifically TNT, RDX, HMX and DNAN; and the frequency of records that studied heavy metals and specifically lead, copper, zinc, arsenic, cadmium and antimony; and the frequency of records studying hydrocarbons. Biological methods were also counted including 16S amplicon sequencing, ITS sequencing, shotgun/metagenomics, nanopore sequencing, phospholipid fatty acid analysis (PLFA), denaturing gradient gel electrophoresis (DGGE), community-level physiological profiling (CLPP), other culture based methods, soil fauna assessments, and the primary soil organism group investigated. Some studies employed multiple biological methods; therefore frequencies represent occurrences rather than mutually exclusive categories. Data analysis and summary statistics were performed using Python-based scripts and spreadsheet analysis.

### 2.7 Methodological Completeness Index

To evaluate the extent to which studies integrated environmental context, contamination assessment, and biological community characterization, a Methodological Completeness Index (MCI) was developed. The index provides a quantitative measure of how comprehensively individual studies reported key components necessary for interpreting soil biological responses in military-impacted environments.

The MCI incorporated four methodological domains: soil physicochemical characterization, contaminant assessment, biological community profiling, and site context reporting. Within each domain, binary variables were assigned to indicate whether specific measurements or descriptors were reported. Soil physicochemical variables included measurements such as soil pH, soil organic carbon or organic matter, total nitrogen, soil texture, soil moisture, electrical conductivity, cation exchange capacity, microbial biomass, and enzyme activity. Contaminant assessment variables included the reporting of explosive compounds (e.g., TNT, RDX, HMX) or other contaminants such as heavy metals or hydrocarbons. Biological characterization variables recorded the use of microbial or soil organism community profiling methods, including sequencing approaches (e.g., 16S rRNA amplicon sequencing, ITS sequencing, shotgun metagenomics), lipid biomarker methods (PLFA), culture-based techniques, and other ecological assessments. Site context variables included descriptors such as site location, military activity type, landscape context, and geographic coordinates.

Scores for each domain were normalized and combined to produce a final MCI score ranging from 0 to 1, where higher values indicate greater methodological completeness. Studies were subsequently categorized into qualitative completeness groups (e.g., very limited, basic, and moderate completeness) to facilitate interpretation of methodological trends across the literature. The index was used to summarize methodological patterns across the included studies and to identify common gaps in environmental characterization within research on military-impacted soils.

## 3 Results

### 3.1 Study selection and screening

A total of 6474 records were identified through database searching across Scopus, Web of Science, Dimensions, PubMed, and Google Scholar. After removal of 85 duplicate records, 6389 studies remained for initial screening. Automated first-stage screening excluded 6327 records based on title, abstract, and keyword relevance criteria, leaving 62 records for second-stage eligibility assessment. Of these, 54 records originated from the structured database search workflow, while 8 additional records were identified through earlier exploratory searching and manual literature identification. Following full-text eligibility assessment, 39 records from the database workflow and 3 records from other sources were excluded. Three studies identified during earlier searching were excluded during eligibility assessment because, although they reported ammunition- or explosive-related contamination, they did not provide sufficient evidence that the contamination originated from a military context. Consequently, 15 studies identified through the systematic search workflow and 5 additional studies identified through prior searching met the inclusion criteria. In total, 20 studies were included in the final review, a table summarising the study characteristics is available below (Table 1). The study selection process followed PRISMA reporting guidelines (Figure 2).

**Table 1.**
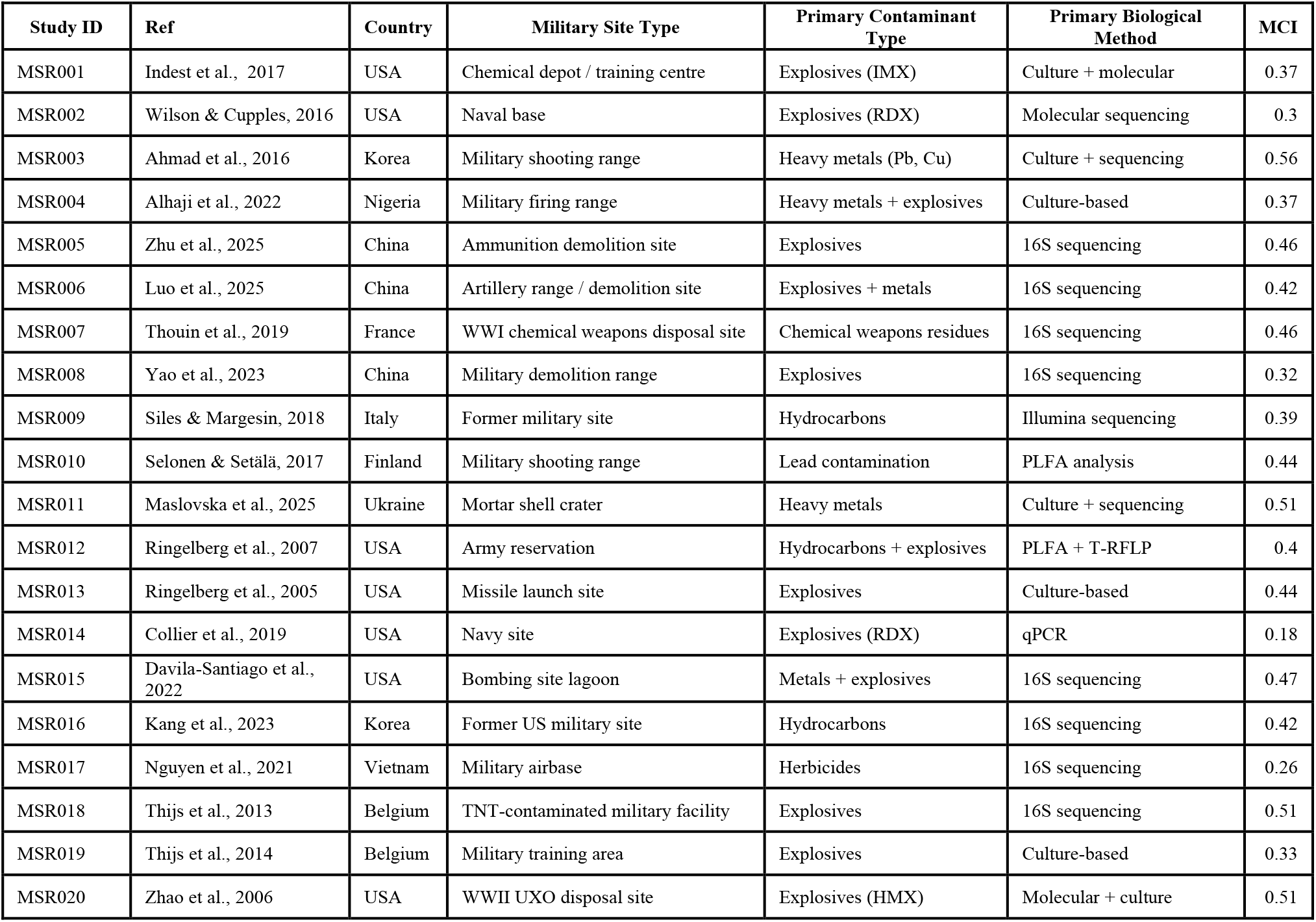
Summary characteristics of included studies.

| Study ID | Ref | Country | Military Site Type | Primary Contaminant Type | Primary Biological Method | MCI |
| --- | --- | --- | --- | --- | --- | --- |
| MSR001 | Indest et al., 2017 | USA | Chemical depot / training centre | Explosives (IMX) | Culture + molecular | 0.37 |
| MSR002 | Wilson & Cupples, 2016 | USA | Naval base | Explosives (RDX) | Molecular sequencing | 0.3 |
| MSR003 | Ahmad et al., 2016 | Korea | Military shooting range | Heavy metals (Pb, Cu) | Culture + sequencing | 0.56 |
| MSR004 | Alhaji et al., 2022 | Nigeria | Military firing range | Heavy metals + explosives | Culture-based | 0.37 |
| MSR005 | Zhu et al., 2025 | China | Ammunition demolition site | Explosives | 16S sequencing | 0.46 |
| MSR006 | Luo et al., 2025 | China | Artillery range / demolition site | Explosives + metals | 16S sequencing | 0.42 |
| MSR007 | Thouin et al., 2019 | France | WWI chemical weapons disposal site | Chemical weapons residues | 16S sequencing | 0.46 |
| MSR008 | Yao et al., 2023 | China | Military demolition range | Explosives | 16S sequencing | 0.32 |
| MSR009 | Siles & Margesin, 2018 | Italy | Former military site | Hydrocarbons | Illumina sequencing | 0.39 |
| MSR010 | Selonen & Setälä, 2017 | Finland | Military shooting range | Lead contamination | PLFA analysis | 0.44 |
| MSR011 | Maslovska et al., 2025 | Ukraine | Mortar shell crater | Heavy metals | Culture + sequencing | 0.51 |
| MSR012 | Ringelberg et al., 2007 | USA | Army reservation | Hydrocarbons + explosives | PLFA + T-RFLP | 0.4 |
| MSR013 | Ringelberg et al., 2005 | USA | Missile launch site | Explosives | Culture-based | 0.44 |
| MSR014 | Collier et al., 2019 | USA | Navy site | Explosives (RDX) | qPCR | 0.18 |
| MSR015 | Davila-Santiago et al., 2022 | USA | Bombing site lagoon | Metals + explosives | 16S sequencing | 0.47 |
| MSR016 | Kang et al., 2023 | Korea | Former US military site | Hydrocarbons | 16S sequencing | 0.42 |
| MSR017 | Nguyen et al., 2021 | Vietnam | Military airbase | Herbicides | 16S sequencing | 0.26 |
| MSR018 | Thijs et al., 2013 | Belgium | TNT-contaminated military facility | Explosives | 16S sequencing | 0.51 |
| MSR019 | Thijs et al., 2014 | Belgium | Military training area | Explosives | Culture-based | 0.33 |
| MSR020 | Zhao et al., 2006 | USA | WWII UXO disposal site | Explosives (HMX) | Molecular + culture | 0.51 |

**Figure 2.**
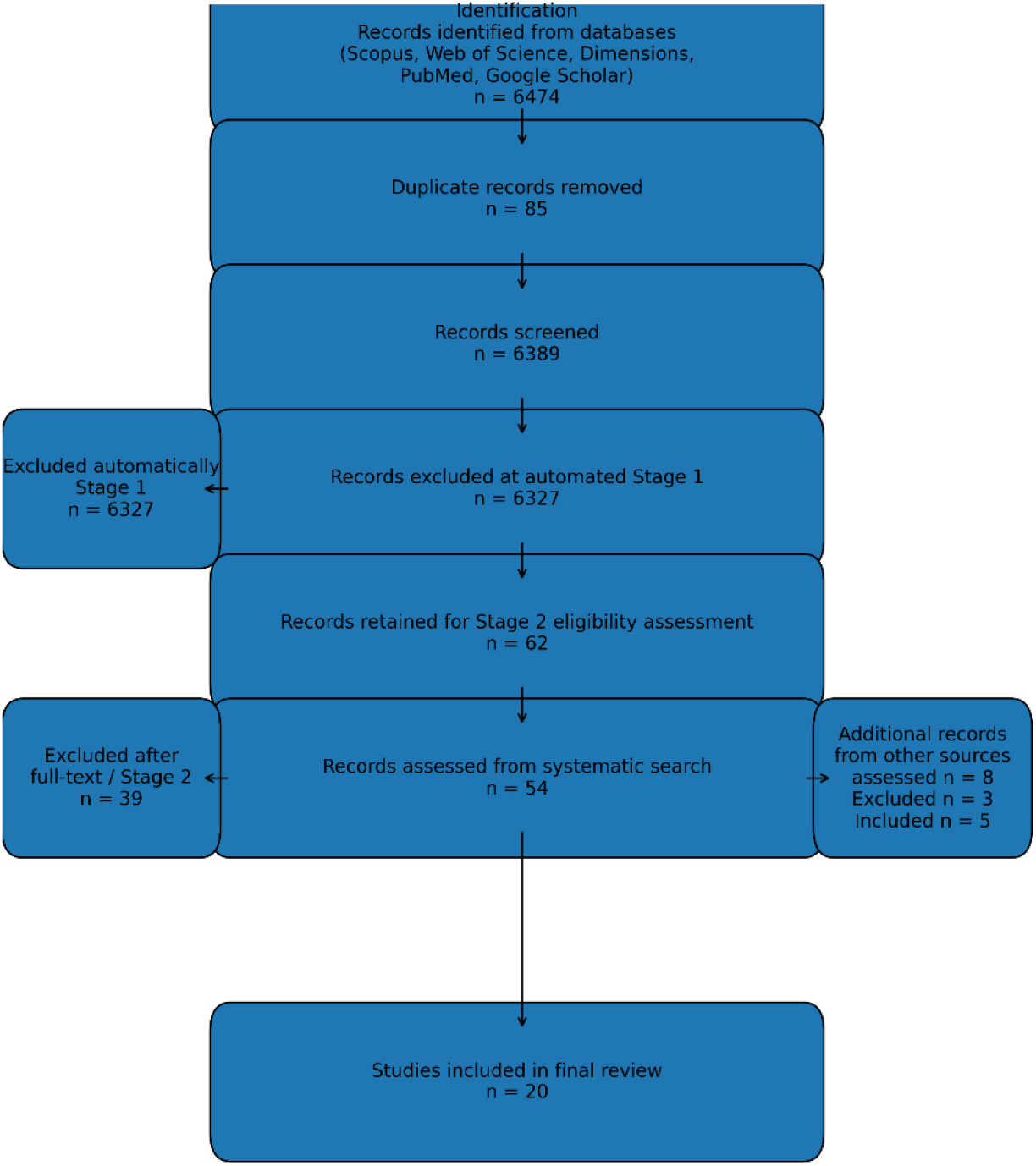
PRISMA flow diagram summarizing the study identification, screening, and eligibility assessment process. A total of 6474 records were identified through database searching. After removal of duplicates and automated screening, 62 studies were assessed for eligibility. Fifteen studies were included from the systematic search workflow and five additional studies were identified from prior searching and manual literature identification, resulting in a final set of 20 studies included in the review.

### 3.2 Soil physicochemical variables measured across studies

The reporting of soil physicochemical properties varied substantially across the included studies (Figure 3). Soil pH was the most frequently reported variable, measured in 12 of the 20 (60 %) studies, followed by soil organic matter and soil texture (45%), which were reported in nine studies each. Soil organic carbon and soil moisture were measured in eight studies (40%), while total nitrogen was reported in 6 studies (30%). Measurements of electrical conductivity (5, 25%) and cation exchange capacity (4, 20%) were less common. Indicators of biological activity within the soil matrix were reported inconsistently; microbial biomass (4, 20%) and enzyme activity (2, 10%). Only one study (5%) reported the soil carbon-to-nitrogen ratio. Notably, none of the included studies reported soil bulk density, a key physical parameter influencing soil structure, compaction, and microbial habitat conditions. Overall, these results indicate that many studies investigating soil organisms in military-impacted environments provide limited characterisation of the physicochemical properties of the soil environment, potentially constraining the interpretation of biological responses to contamination and disturbance. This uneven reporting of soil properties highlights a methodological gap in the literature, where biological community analyses are frequently conducted without comprehensive environmental context.

**Figure 3.**
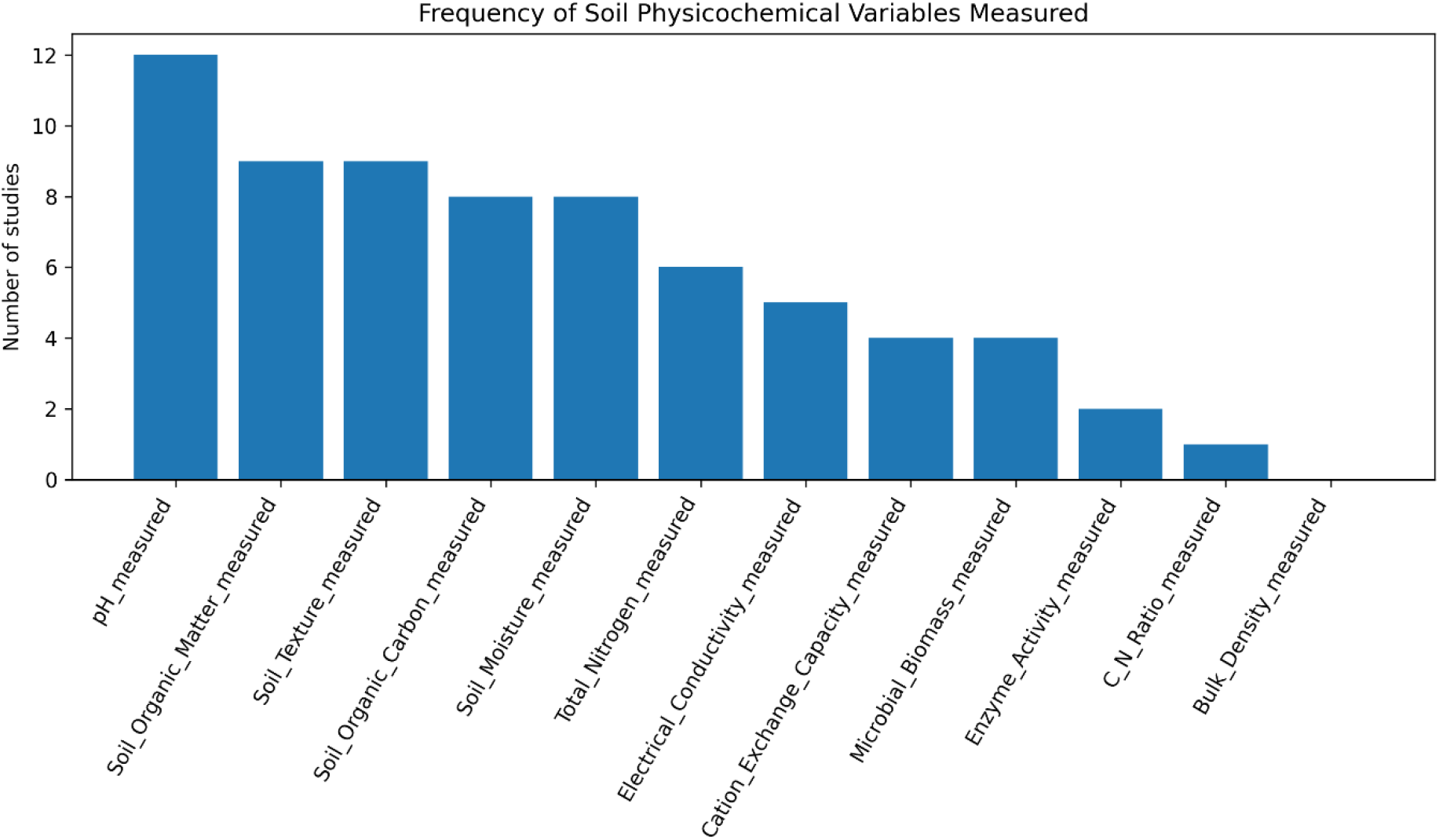
Frequency of soil physicochemical variables measured across the studies included in this review (n = 20). The figure illustrates the number of studies reporting measurements of key soil properties, including pH, soil organic matter (SOM), soil organic carbon (SOC), soil texture, soil moisture, total nitrogen (TN), electrical conductivity (EC), cation exchange capacity (CEC), microbial biomass, enzyme activity, and the carbon-to-nitrogen (C:N) ratio. Bulk density was not reported in any of the included studies.

### 3.3 Biological methods used for community profiling

A range of approaches were used to characterise soil biological communities across the included studies, although sequencing-based bacterial community analysis dominated the literature (Figure 4). 16S rRNA amplicon sequencing was the most commonly used method, reported in 16 of the 20 studies, reflecting the widespread adoption of high-throughput sequencing approaches for bacterial community profiling. Culture-based methods were also frequently employed, appearing in 12 studies, sometimes to isolate contaminant-tolerant or degradation-capable bacterial strains. Less frequently used methods included phospholipid fatty acid (PLFA) analysis, which was reported in four studies, and shotgun metagenomic sequencing, which was used in two studies to investigate microbial functional potential. Fungal community profiling was rarely reported, with ITS sequencing used in only one study, while soil fauna assessments were also reported in only a single study. No studies in the dataset reported the use of Nanopore sequencing, Biolog community-level physiological profiling, or older molecular fingerprinting techniques such as DGGE, T-RFLP, or ARISA.

**Figure 4.**
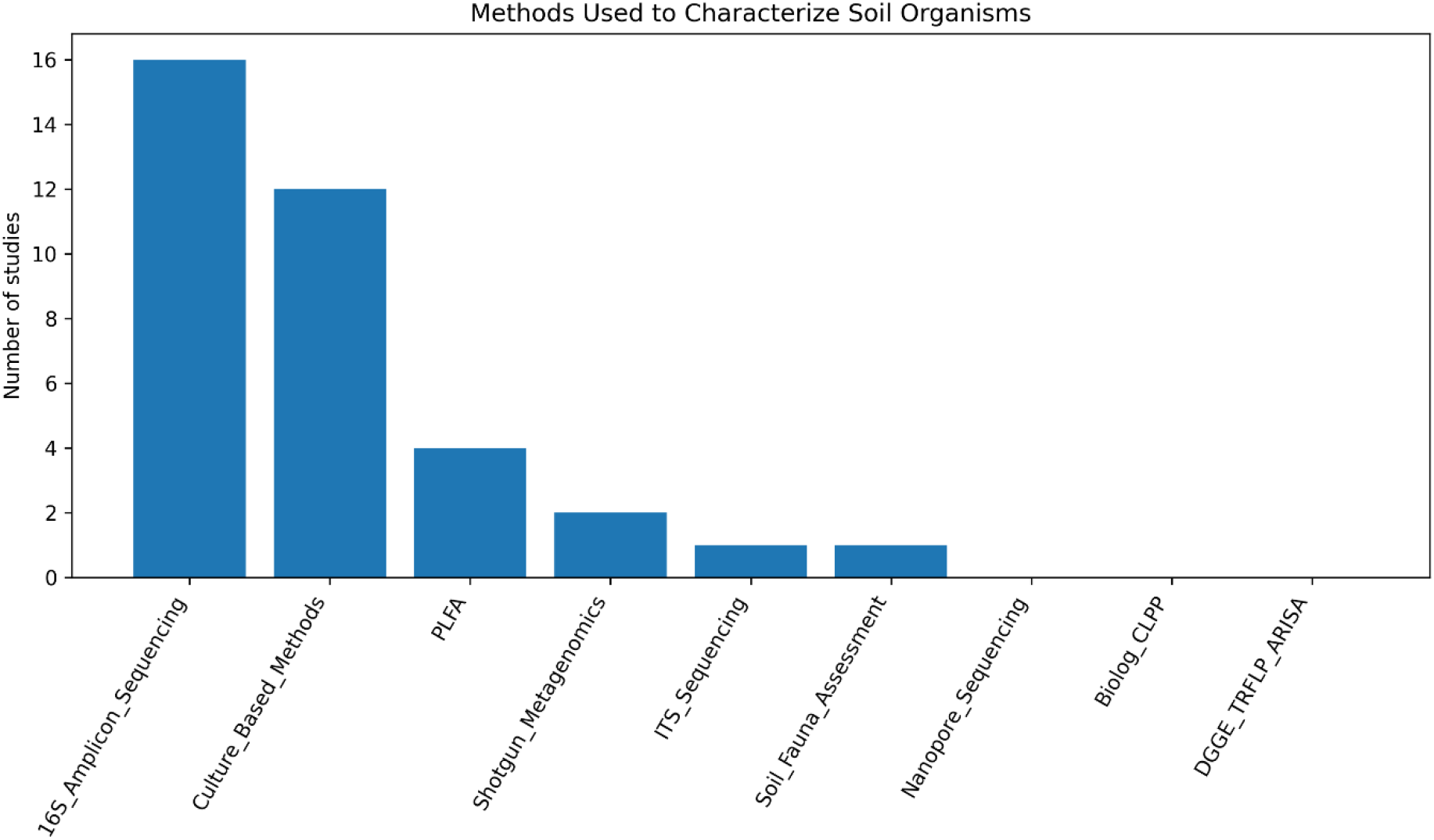
Frequency of biological methods used to investigate soil organisms in the included studies (n = 20). Bars represent the number of studies reporting each method. Because some studies used multiple biological methods, counts represent occurrence frequencies rather than mutually exclusive categories. The most frequently applied technique was 16S rRNA gene amplicon sequencing for bacterial community profiling. Other approaches included culture-based microbial isolation, phospholipid fatty acid (PLFA) analysis, shotgun metagenomics, ITS sequencing for fungal communities, and soil fauna assessment. Some studies employed multiple methods; therefore, frequencies represent occurrences rather than mutually exclusive categories.

Several studies employed multiple biological methods within the same investigation, most commonly combining 16S amplicon sequencing with culture-based isolation techniques, or combining PLFA with culture-based approaches. Consequently, the frequencies reported here represent occurrence frequencies rather than mutually exclusive categories. Overall, these results indicate a strong methodological emphasis on bacterial community characterisation, with comparatively limited attention to fungal communities, soil fauna, or broader ecological community interactions in military-impacted soils.

### 3.4 Contaminant types investigated in military-impacted soils

The studies included in this review investigated a range of contaminants associated with military activities, with a strong emphasis on explosive compounds and heavy metals (Figure 5). Explosive residues were reported in nine of the 20 studies, with the most frequently identified compounds including RDX (reported in six studies), TNT (four studies), and HMX (three studies). Newer or less commonly studied energetic compounds such as DNAN were reported in only one study. Heavy metals were investigated in eight studies, with lead (Pb) being the most commonly reported metal (seven studies), followed by copper (Cu) and arsenic (As) (five studies each), zinc (Zn) and cadmium (Cd) (four studies each), and antimony (Sb) (one study). In addition to explosive compounds and metals, hydrocarbon contamination was reported in four studies, typically associated with fuel storage, vehicle operation, or historical military infrastructure. Several studies investigated multiple contaminant types within the same site, reflecting the complex contamination profiles often associated with military training areas, ammunition disposal sites, and historical conflict locations. Overall, the contaminant profiles reported across the literature indicate that research on military-impacted soils has largely focused on explosive residues and heavy metal contamination, with comparatively fewer studies addressing petroleum hydrocarbons or emerging contaminants.

**Figure 5.**
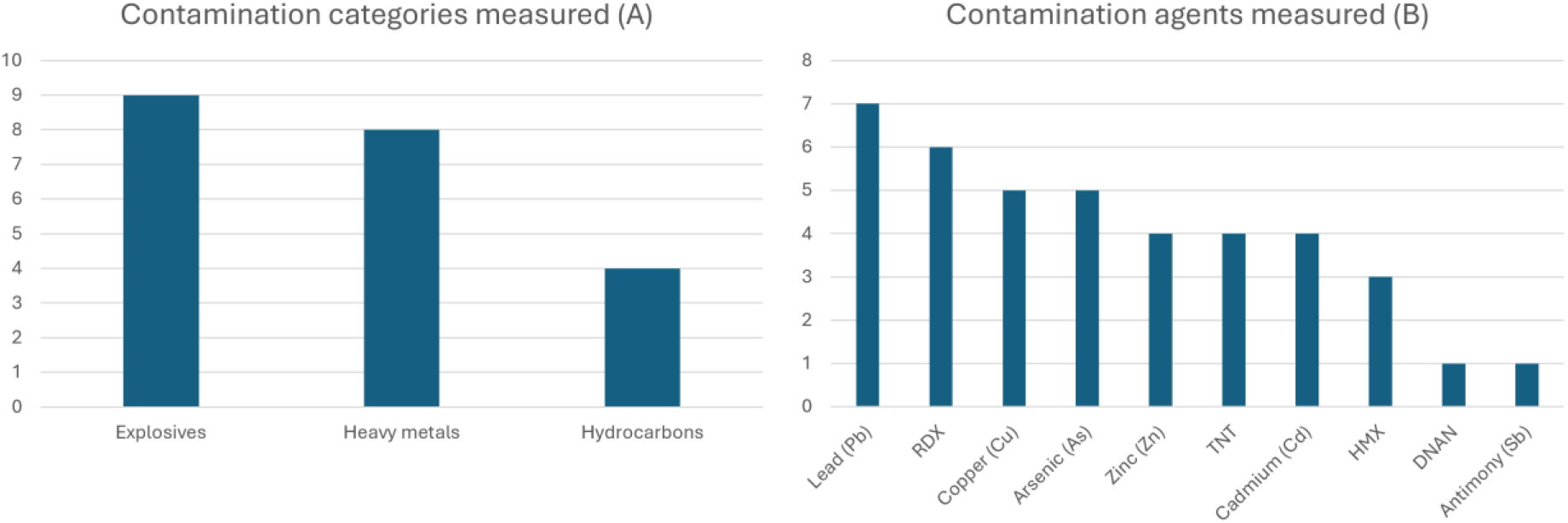
Contamination types investigated across the included studies. The figure shows the number of studies reporting different contaminant categories (A) (explosives, heavy metals and hydrocarbons) and (B) specific contaminants (including explosive compounds (TNT, RDX, HMX), heavy metals (lead, copper, zinc, arsenic, cadmium). Some studies reported multiple contaminants, therefore counts represent occurrence frequencies rather than mutually exclusive categories. Explosive compounds were investigated in 9 of 20 studies (45%), while heavy metals were reported in 8 studies (40%), and hydrocarbons in 4 studies (20%). Among explosive compounds, RDX was the most frequently reported (6 studies), followed by TNT (4 studies) and HMX (3 studies). Explosive compounds and heavy metals were the most commonly investigated contaminants in military-impacted soils. Heavy metal contamination was most commonly associated with lead (Pb), reported in 7 studies, reflecting the influence of small-arms ammunition residues at military training ranges. Other frequently reported metals included copper (Cu) and arsenic (As) (5 studies each), zinc (Zn) and cadmium (Cd) (4 studies each). Emerging insensitive munitions compounds such as 2,4-dinitroanisole (DNAN) were rarely investigated, appearing in only one study.

### 3.5 Geographic distribution of studies

The geographic distribution of studies investigating soil organisms in military-impacted environments was geographically diverse but unevenly distributed among a limited number of countries (Figure 6). The United States accounted for the largest number of studies (7 of 20; 35%), reflecting extensive research conducted at military installations and contaminated training areas. China contributed three studies (15%), while Belgium contributed two studies (10%), largely associated with historical military contamination and remediation research. The remaining studies were distributed across several countries, including Korea, Nigeria, France, Italy, Finland, Ukraine, Puerto Rico (USA), the Republic of Korea, and Vietnam, each represented by a single study. Several of these sites corresponded to historically contaminated military locations such as training ranges, ammunition demolition sites, airbases, and former conflict zones. Overall, the results indicate that research on soil biological communities in military-impacted environments is geographically concentrated in a small number of countries, with large regions of the world remaining poorly represented in the literature, particularly areas affected by historical or ongoing armed conflict.

**Figure 6.**
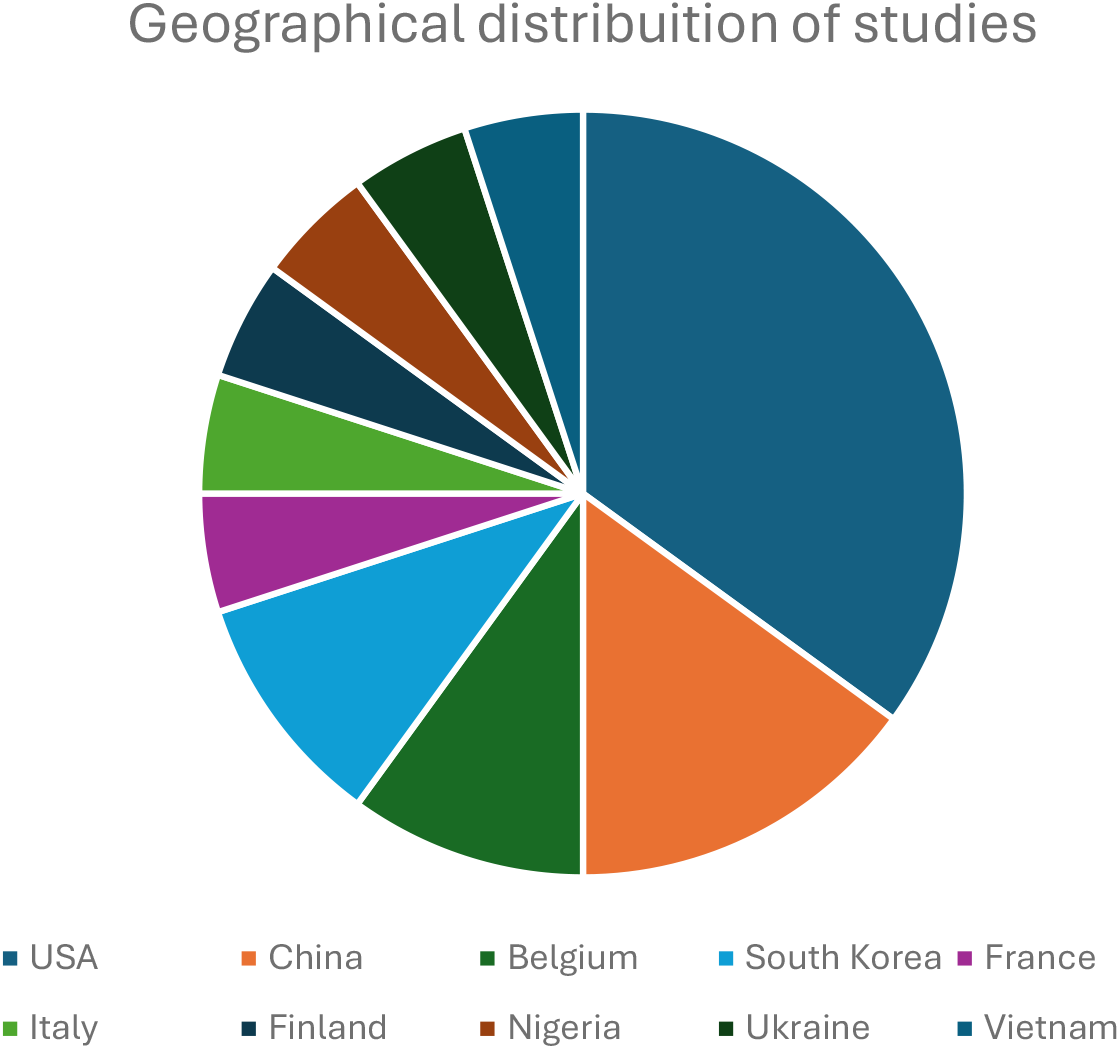
Geographic distribution of studies investigating soil organisms in military-impacted environments. The figure shows the number of studies conducted in each country represented in the review (n = 20). Research efforts were concentrated in a limited number of countries, particularly the United States and China, with fewer studies conducted in regions directly affected by recent or ongoing armed conflict. Notably, relatively few studies were conducted in regions experiencing recent or ongoing armed conflict, despite the potential for substantial soil disturbance and contamination in such environments.

### 3.6 Methodological completeness of studies

The methodological completeness of the included studies varied considerably, as reflected by the Methodological Completeness Index (MCI) scores calculated for each study (Figure 7). Across the dataset, MCI values ranged from 0.175 to 0.561, with a mean score of 0.405 and a median score of 0.421, indicating that most studies incorporated only a subset of the environmental and methodological components considered important for interpreting biological responses in contaminated soils. The majority of studies fell within the basic methodology category, reflecting limited integration of soil physicochemical characterisation, contaminant assessment, and biological community profiling. A smaller number of studies achieved moderate methodological completeness, typically through the combined reporting of soil properties, contaminant concentrations, and microbial community sequencing. Only a small number of studies provided comprehensive environmental characterisation alongside biological analysis. Overall, the results suggest that research investigating soil organisms in military-impacted environments frequently emphasises microbial community characterisation while providing comparatively limited contextual information on soil properties and environmental conditions, which may constrain interpretation of ecological responses to contamination and disturbance.

**Figure 7.**
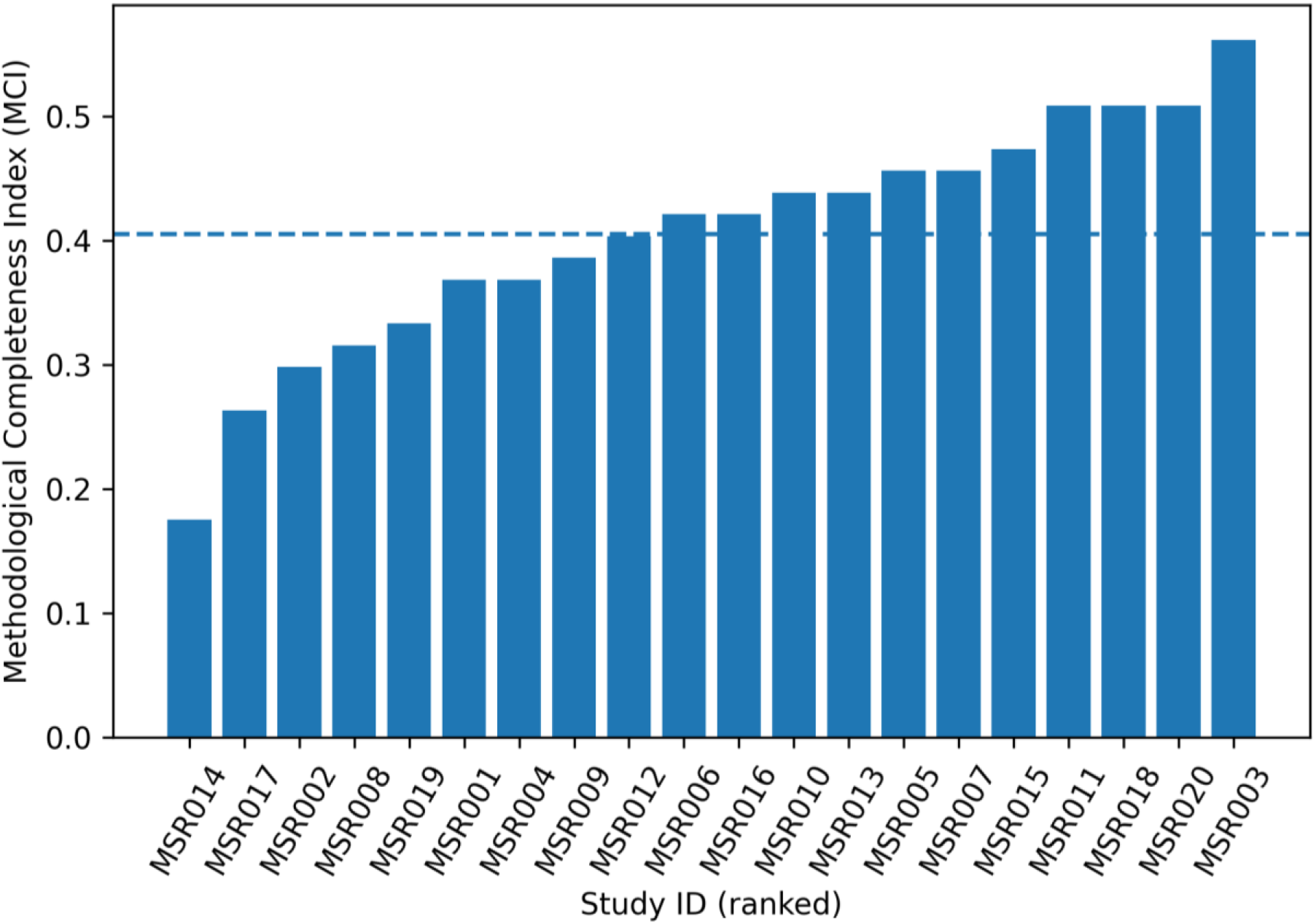
Methodological completeness of included studies. Methodological Completeness Index (MCI) scores for the included studies (n = 20), ranked from lowest to highest. The index integrates reporting of soil physicochemical variables, contaminant characterisation, biological community profiling methods, and site context information. The dashed horizontal line indicates the mean MCI value across all studies (0.405).

## 4 Discussion

### 4.1 Overview of key findings

This systematic review synthesised available studies investigating soil microbial communities in military-impacted environments using sequencing-based and related biological methods. Across the included studies, several clear patterns emerged. First, the literature is dominated by studies investigating bacterial communities using 16S rRNA amplicon sequencing, while fungal communities and soil fauna are rarely examined. Second, soil physicochemical characterization was highly variable across studies, with many investigations measuring only a limited set of environmental parameters. Third, research has primarily focused on legacy explosive compounds and heavy metals, with comparatively few studies examining other contamination types associated with military activities. Finally, the geographic distribution of studies reveals strong concentration in a small number of countries, with limited representation from regions currently affected by armed conflict.

Military-impacted soils are frequently studied without explicit reference to military activity in titles or abstracts. As a result, systematic searches relying solely on military keywords may miss relevant studies, requiring broader environmental searches combined with manual screening.

Together, these findings highlight both the progress made in applying molecular tools to military soil ecosystems and the substantial methodological and geographic gaps that remain.

- Military soil microbiology studies lack comprehensive soil characterization
- The literature focuses heavily on legacy explosive contamination
- 16S sequencing dominates, while fungal and soil fauna studies are rare
- Major geographic gaps exist in conflict-affected regions

### 4.2 Methodological approaches used to study soil organisms

The analysis revealed a strong reliance on bacterial community profiling using 16S rRNA gene amplicon sequencing, which was employed in the majority of studies included in this review. The widespread adoption of 16S sequencing reflects its accessibility, cost-effectiveness, and ability to provide broad insights into bacterial community composition (Smets et al., 2016; Jimenez, 2025; Ceretto & Weinig, 2025). However, this methodological focus also highlights important limitations in current research. Firstly, the full 16S region may provide better taxonomic classification (Bertolo et al., 2024). Secondly, fungal community analysis was rarely performed, despite fungi playing critical roles in soil nutrient cycling and ecosystem resilience (Frąc et al., 2028; Martínez-García et al., 2023). Similarly, only a single study assessed soil fauna, even though organisms such as nematodes, microarthropods, and earthworms are important indicators of soil ecological condition. Several studies combined molecular methods with culture-based approaches, suggesting that traditional microbiological techniques remain valuable for isolating organisms capable of degrading explosive compounds or tolerating heavy metals. However, relatively few studies employed more advanced approaches such as shotgun metagenomics, which can provide deeper insight into functional potential and metabolic pathways. Emerging sequencing platforms, including nanopore sequencing technologies, were not represented in the included studies, indicating that their application to military soil ecosystems remains limited.

### 4.3 Soil physicochemical characterisation in military soil studies

The results of this review demonstrate considerable variability in the measurement of soil physicochemical variables across studies. While basic parameters such as soil pH, soil organic matter, and soil organic carbon were frequently reported, many other important indicators of soil function were measured in relatively few studies. For example, microbial biomass and enzyme activity – important indicators of microbial functional activity – were reported in only a small proportion of studies. Notably, none of the included studies reported measurements of soil bulk density, despite the well-documented influence of military activities such as vehicle movement, artillery fire, and troop movement on soil compaction and physical structure. The limited measurement of soil physical and chemical parameters may constrain the interpretation of microbial community data. Soil microbial communities are strongly influenced by environmental variables including moisture, nutrient availability, soil texture, and structural characteristics. Without comprehensive environmental characterisation, it becomes difficult to determine whether observed microbial community changes are driven by contamination, soil disturbance, or natural variation in soil properties.

### 4.4 Contamination types investigated in military soils

Contamination patterns observed across the reviewed studies indicate a strong research focus on explosive compounds and heavy metals, particularly those associated with ammunition residues and training range activities. Explosive compounds such as TNT, RDX, and HMX were among the most frequently reported contaminants, reflecting historical concerns about explosive contamination at military training ranges and ammunition disposal sites. Heavy metals, particularly lead, were also widely reported, likely due to their association with small-arms ammunition used at shooting ranges.

In contrast, other contamination types commonly associated with military activities were investigated less frequently. For example, hydrocarbon contamination, which can arise from fuel spills, vehicle operations, and aircraft maintenance activities, was examined in relatively few studies. Similarly, newer insensitive munitions compounds, which are increasingly used in modern military applications, were rarely investigated. This suggests that research has primarily focused on legacy contaminants rather than emerging contaminants associated with modern military technologies.

The contamination patterns identified in this review suggest a strong research focus on legacy explosive compounds and lead-based ammunition residues. Compounds such as TNT, RDX, and HMX were frequently investigated, reflecting historical contamination concerns at military training ranges and ammunition disposal sites. In contrast, modern insensitive munitions compounds such as DNAN were rarely examined. This is notable given the increasing adoption of insensitive munitions by modern armed forces to improve handling safety and reduce accidental detonations. Similarly, hydrocarbons associated with fuel spills and vehicle operations were investigated in relatively few studies. These findings highlight an important research gap in understanding the broader range of contaminants associated with military activities.

### 4.5 Geographic distribution of research

The geographic distribution of studies included in this review reveals a concentration of research activity in a small number of countries. The United States accounted for the largest proportion of studies, followed by China and several European countries. This pattern likely reflects the availability of research funding, environmental monitoring programs, and established research infrastructure in these regions. However, the distribution also highlights an important gap in the current literature. Only a small number of studies were conducted in regions that have experienced recent or ongoing armed conflict. This may reflect the practical and logistical challenges associated with conducting environmental research in conflict-affected landscapes, including issues related to safety, accessibility, and funding availability. Nevertheless, the limited number of studies from these regions suggests that the ecological consequences of military activity in many parts of the world remain poorly understood.

### 4.6 Methodological completeness and limitations of existing studies

The methodological completeness analysis conducted in this review provides further insight into the limitations of existing studies. The mean methodological completeness index (MCI) was relatively low, indicating that most studies incorporated only a subset of the methodological components considered important for comprehensive soil ecosystem assessment. Many studies focused primarily on microbial community sequencing while providing limited information about soil physicochemical properties, contamination concentrations, or site-level environmental context. This limited methodological integration may reduce the ability to identify causal relationships between contamination and biological responses. Microbial community structure is shaped by complex interactions among environmental conditions, contaminant concentrations, and ecological processes. Studies that measure only microbial composition without detailed environmental characterization may therefore provide an incomplete picture of ecosystem responses to military disturbance. This limits the ability to link microbial community patterns to environmental drivers and reduces comparability across studies. Future research should prioritize integrated study designs that combine microbial community profiling with detailed measurements of soil physicochemical properties, contamination concentrations, and landscape disturbance indicators.

### 4.7 Recommendations for future research

The findings of this review suggest several priorities for future research on military-impacted soils. First, studies should adopt more integrated methodological approaches that combine microbial community analysis with detailed measurements of soil physicochemical properties and contaminant concentrations. Such integrated approaches would enable stronger inference about the environmental drivers of microbial community change. Second, greater attention should be given to understudied biological groups, including fungi and soil fauna, which play key roles in soil ecosystem functioning. Expanding the taxonomic scope of studies would provide a more comprehensive understanding of ecological responses to military disturbance. Third, research should increasingly address emerging contaminants, including compounds associated with insensitive munitions and other modern military materials. Finally, there is a need for more studies conducted in regions affected by recent or ongoing armed conflict, where the ecological consequences of military activities may be particularly significant.

### 4.8 Limitations of the review

Several limitations should be considered when interpreting the results of this review. Although multiple databases were searched using structured search strategies, it is possible that some relevant studies were not captured, particularly those published in non-English languages or in grey literature sources. In addition, variation in reporting practices among studies limited the ability to extract consistent information on certain soil properties and contamination characteristics. The methodological completeness index used in this study provides a useful comparative metric but should be interpreted as an approximate indicator of methodological coverage rather than a definitive measure of study quality.

Overall, this systematic review demonstrates that research on soil microbial communities in military-impacted environments remains relatively limited and methodologically heterogeneous. While advances in sequencing technologies have enabled detailed characterization of bacterial communities in contaminated soils, many studies lack comprehensive environmental context and focus primarily on a narrow range of contaminants. Addressing these gaps will require more integrated study designs, broader geographic coverage, and expanded investigation of diverse soil organisms. Such efforts will be essential for improving understanding of the ecological consequences of military activities and for informing the management and restoration of military-impacted landscapes.

## Notes

### Competing Interest Statement

The authors have declared no competing interest.

